# On enhancing variation detection through pan-genome indexing

**DOI:** 10.1101/021444

**Authors:** Daniel Valenzuela, Niko Välimäki, Esa Pitkänen, Veli Mäkinen

**Affiliations:** Helsinki Institute for Information Technology HIIT, Department of Computer Science, University of Helsinki, P. O. Box 68 (Gustaf Hällströmin katu 2b), Helsinki FIN-00014, Finland; Genome-Scale Biology Research Program, Research Programs Unit, University of Helsinki, Biomedicum, P. O. Box 63 (Haartmaninkatu 8), Helsinki FIN-00014, Finland; Department of Medical Genetics, Biomedicum, University of Helsinki, Finland

## Abstract

Detection of genomic variants is commonly conducted by aligning a set of reads sequenced from an individual to the reference genome of the species and analyzing the resulting *read pileup*. Typically, this process finds a subset of variants already reported in databases and additional novel variants characteristic to the sequenced individual. Most of the effort in the literature has been put to the alignment problem on a single reference sequence, although our gathered knowledge on species such as human is *pan-genomic*: We know most of the common variation in addition to the reference sequence. There have been some efforts to exploit *pan-genome indexing*, where the most widely adopted approach is to build an index structure on a set of reference sequences containing observed variation combinations.

The enhancement in alignment accuracy when using pan-genome indexing has been demonstrated in experiments, but so far the above *multiple references* pan-genome indexing approach has not that much been tested on its final goal, that is, in enhancing variation detection. This is the focus of this article: We study a generic approach to add variation detection support on top of the multiple references pan-genomic indexing approach. Namely, we study the read pileup on a multiple alignment of reference genomes, and propose a heaviest path algorithm to extract a new recombined reference sequence. This recombined reference sequence can then be utilized in any standard read alignment and variation detection workflow. We demonstrate that the approach enhances variation detection on realistic data sets.

## Introduction

With the new technologies for high-throughput sequencing, we have witnessed the emergence of enormous genomic databases. While detailed collections of human genetic variation are growing, the algorithmic community is struggling to offer proper techniques to manage these volumes of information efficiently.

An important breakthrough occurred when the Burrows-Wheeler Transform^1^ greatly improved the state of the art of indexed pattern matching.^2, 3^ Years later, based on those techniques, we witnessed the rise of many read aligners like the Burrows-Wheeler Aligner (BWA),^4^ Bowtie^5^ and SOAP,^6^ among many others.

While standard aligners work by indexing one reference sequence, and then searching for the best alignment of the reads, we are interested in a more recent approach, namely, *pan-genomic indexing*.^7–16^ That is, instead of having one reference sequence, an entire collection is indexed, allowing the reads to be mapped against any genome of the reference set or even to some recombination of them. We will explore new possibilities to use pan-genome indexing for variation calling.

*Variation calling* is the process of identifying the differences in the genome of an individual from the consensus of the population. The standard approach is to obtain a set of reads from the donor individual, and align them against the consensus. The resulting *read pileup* is further analyzed to discover the variants, see e.g.^17, 18^

Since an individual contains a unique combination of variations common in the population and some unique novel variations, the set of called variants is largely overlapping with a database of variations gathered in earlier studies. One way to exploit this gathered genetic knowledge is to build a *population automaton* recognizing all variation combinations.^13^ In this approach, a population automaton is indexed with an extension of the Burrows-Wheeler transform to support efficient read alignment. Experiments on variation-rich regions of human genome show that the read alignment accuracy is greatly improved over the standard approach.

The caveat of the population automaton approach to pan-genome indexing is the indexing phase: The worst case index size is exponential, so typically some variants need to be dropped from the automaton with a manual trial-and-error approach to guarantee the good expected case behaviour.^13^ Alternatively, one can enumerate all close-by variant combinations and index the resulting variant contexts (i.e. short subpaths in population automaton) in addition to the reference.^7, 10, 14, 15^ Yet, in these approaches, one always has to restrict to short context length in order to avoid exponential blowup.

Due to the complexity of the population automata / context enumeration approaches, a majority of the literature on pan-genome indexing is focusing on the more feasible input of a set of individual genomes.^8, 9, 11, 12, 16^ For such input, the Burrows-Wheeler transform is of linear size, and the shared content among individuals results into highly compressed indexes that take space close to size of the index of a single reference sequence. Lately, there have been proposals to use Lempel-Ziv indexing to obtain extremely well compressed indexed that still support efficient read alignment.^9, 11, 12, 16^

The literature on the above multiple references pan-genome indexing has largely concentrated on the time and space efficiency aspects, neglecting its final goal of enhancing variation calling. This article aims to fill this gap.

We study a generic approach to add variation detection support on top of the multiple references pan-genomic indexing approach. Namely, we study the read pileup on a multiple alignment of reference genomes, and propose a heaviest path algorithm to extract a new recombined reference sequence. This recombined *ad hoc* reference sequence can then be utilized in any standard read alignment and variation detection workflow. We demonstrate that the approach enhances variation detection on realistic data sets. With the graph-based approach such improvement has already been reported in the seminal paper^7^ and very recently in a detailed study of MHC region on human chromosome 6.^15^ Our generic approach adds the possibility to use scalable multiple reference pan-genome indexes for the same purposes.

Our overall scheme is illustrated in Fig. 1.

**Figure 1.**
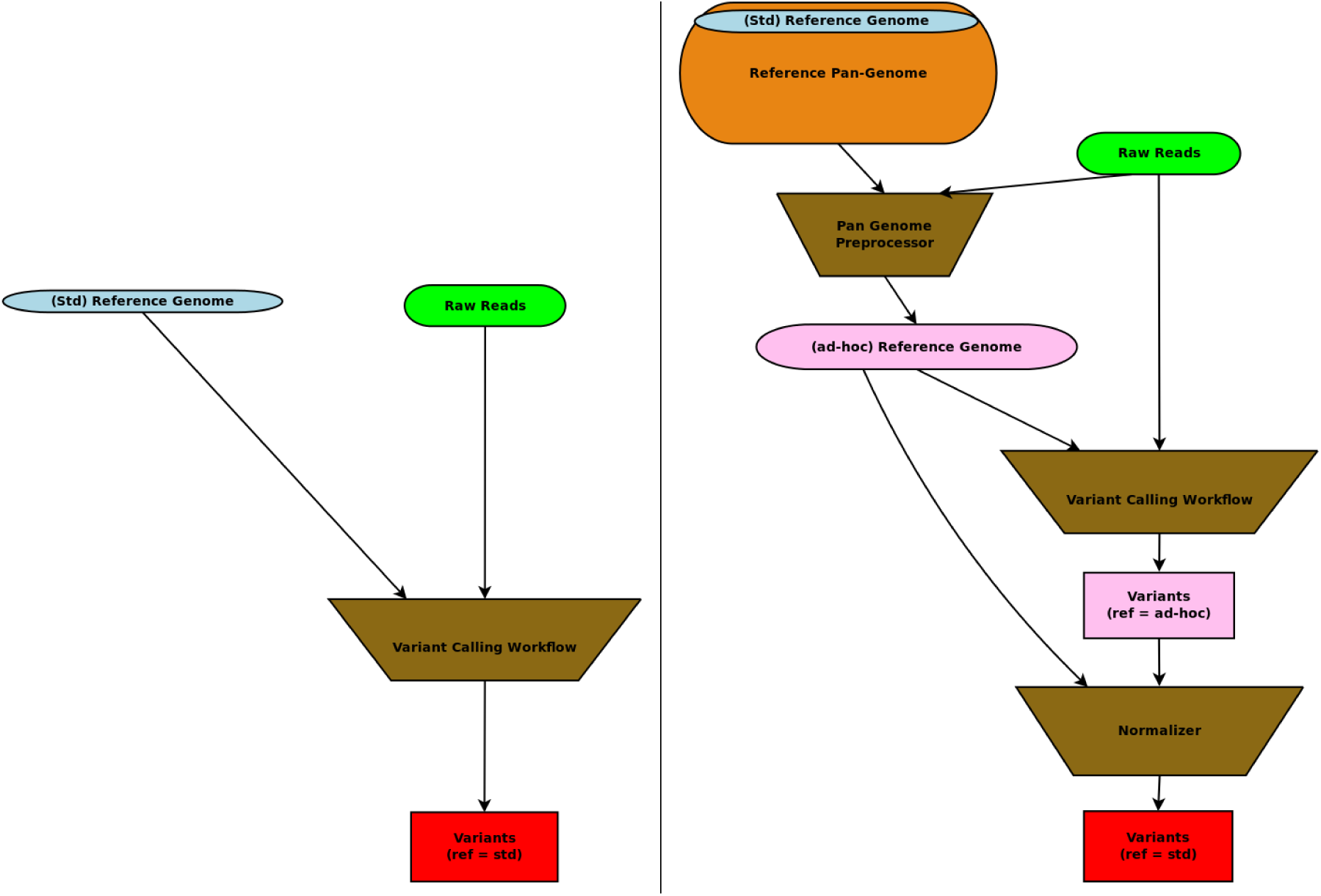
Left: schematic view of a variant calling workflow using a standard reference. Right: Our pan-genomic workflow, where we use the traditional workflow as a subcomponent.

## Results

The components of our scheme are detailed in the Methods section, but for the evaluation of the results it is enough to understand the final goal of predicting the donor genome more accurately than with the standard approach.

### Experimental setup and evaluation measures

Our experimental setup consists of a hidden donor genome, out of which a set of sequencing reads is given as input to the variation calling prediction workflows. The standard approach we compare against, is the GATK best practices workflow^17^ that aligns the reads against a reference genome and analyses the resulting read pileup. Our scheme aligns the reads against multiple reference genomes and generates an *ad hoc* reference which is given to GATK best practices workflow in place of the standard reference. We also compare against the pan-genome reference graph approach,^13^ which we modified also to output an *ad hoc* reference (see Methods section), so that one can apply the same GATK best practices workflow also for that.

All evaluated methods output variation calling results that are projected with respect to the standard reference genome. Our hidden donor genome can also be represented as a set of variants with respect to the standard reference genome. This means that we can calculate the standard prediction success measures such as precision and recall. For this, we chose to define the prediction events per base, rather than per variant, to tolerate better invariances of variant locations as found critical in a recent study.^19^ We separate the single nucleotide variant (SNV) calls from indel calls as the results differ clearly for these two subclasses. A true positive (TP) SNV call is a SNV in the true donor and in the predicted donor. A false positive (FP) SNV call is not a SNV in the true donor but is a SNV in the predicted donor. A false negative (FN) SNV call is a SNV in the true donor but is not a SNV in the predicted donor. A true positive (TP) indel call is either an inserted base in the true donor with an identical inserted base in the predicted donor, or a deleted base in both the true and predicted donor. A false positive (FP) indel call is neither inserted nor deleted base in the true donor but is either inserted or deleted base in the predicted donor. A false negative (FN) indel call is an inserted or deleted base in the true donor but is neither inserted nor deleted base in the predicted donor. We report *precision*=TP/(TP+FP) and *recall*=TP/(TP+FN).

In addition to precision and recall, we also compute the unit cost edit distance of the true donor and the predicted donor. This is defined as the minimum amount of single base substitutions, insertions, and deletions required to convert the predicted donor into the true donor. Here the sequence content of the true donor is constructed by applying its set of variants to the standard reference and the sequence content of the predicted donor is constructed by applying the predicted variants either to the standard reference (GATK) or to the *ad hoc* reference (our methods).

There are good incentives to use this measure to complement precision and recall: First, it gives one number reflecting how close the prediction is to the ground truth. Second, the projection from the *ad hoc* reference to the standard reference may loose information. Last, repeat- and error-aware direct comparison of indel variant predictions is non-trivial and only handled properly on deletions.^19^

As our experiments are on human data, where genomes are diploids, the heterozygous variants may overlap, which causes some changes to the evaluation measures above. For our evaluation purposes, we treat all variants as they were homozygous. That is, when applying the variants to the reference, we omit variants that overlap already processed ones, and the result is thus a single sequence consisting of all compatible variants. We follow this approach also when computing the precision and recall measures to make the “per base” prediction events well-defined.

### Experiment on highly-polymorphic regions

Graph-based pan-genome indexing has been thoroughly evaluated in^13^ for mapping accuracy on human genome data. From those results one can infer the common sense result that on areas with isolated short indels and SNVs, a regular single-reference based indexing approach with a highly engineered alignment algorithm is already sufficient. Hence, the focus of pan-genome indexing for read alignment and variation calling should be on complex regions with larger indels and/or densely located simpler variants, where significant improvements are still possible. For this goal, a test setup based on the analysis of highly-polymorphic regions of the human genome^20, 21^ was created in.^13^ We followed this test setup consisting of variation-rich regions from 93 genotyped Finnish individuals (1000 genomes project, phase 1 data). The 93 diploid genomes gave us a multiple alignment of 186 strains plus the GRCh37 consensus reference. We chose variation-rich regions that had 10 SNVs within 200 bases or less. The total length of these regions was 2.2 MB. To produce the ground-truth data for our experimental setup, we generated 221559 100 bp single-end reads from each of the Finnish individuals giving an average coverage of 10*x*.

To validate our hypothesis that having a richer set of genomes as a reference will improve the quality of the variation calling process we tested our preprocessor scheme using a reference set of the first 20, 50, and 100 genomes of the 186. The genomes 101 to 186 are the potential donors. The results are illustrated in Tables 1 and 2. Row GATK of Table 1 stands for the GATK Bests Practices workflow. Row MSA + GATK of Table 1 stands for the multiple sequence alignment -based pan-genome indexing scheme specified in the Methods section to produce the *ad hoc* reference. Row Graph +GATK of Table 1 is using the graph-based indexing of,^13^ where the *ad hoc* reference component is specified in the Methods section.

**Table 1.** Edit distance from the predicted donor sequence to the true donor. The average distance between the true donors and the reference is 95193,9.

|  | Pan genome reference size |  |  |  |
| --- | --- | --- | --- | --- |
|  | 1 | 20 | 50 | 100 |
| GATK | 74695,9 | - | - | - |
| MSA + GATK | - | 2885,5 | 1956,9 | 1204,7 |
| Graph +GATK | - | 3230,4 | 3336,8 | 2706,9 |

**Table 2.** Precision and recall of our method MSA compared to GATK.

| Measure | GATK | 20 | 50 | 100 |
| --- | --- | --- | --- | --- |
| SNV Precision | 0.992161 | 0.997355 | 0.996913 | 0.996348 |
| SNV Recall | 0.904897 | 0.996721 | 0.997528 | 0.997554 |
| Indel Precision | 0.364853 | 0.994608 | 0.994906 | 0.994804 |
| Indel Recall | 0.0624981 | 0.961927 | 0.973369 | 0.982674 |

The columns in both tables refer to predictions after using GATK with standard reference, and after using *ad hoc* references produced using different size sets of reference genomes in pan-genome indexing.

The results are averages over 10 donors.

## Discussion

Our results indicate that using pan-genome indexing indeed improves variation calling results significantly on highly-polymorphic regions of human genome: Edit distance of predicted donor and true donor is much smaller already in the case when 10 references are used in place of one, and the distance gets smaller the more references are used. The same behaviour can be observed from the precision and recall results, where especially indel calls are improved significantly after the use of pan-genome indexing. These results reconfirm the findings in^7, 15^ about the graph-based approach to pan-genome indexing.

A unique feature of our proposal is the genericity; the approach works both on graph representations and on multiple alignment representations of a pan-genome. Earlier studies on pan-genome indexing have mostly focused on read alignments, which are then normalized to the reference for compatibility with the existing variant calling workflows. Here we proposed instead to globally analyse all read alignments and to produce an *ad hoc* reference that can be used in place of the standard reference. We keep the projection between the *ad hoc* reference and the standard reference, so that the variation calling results can always be normalized to the standard reference afterwards.

One aspect we did not study here is the scalability of the approach – an aspect that is however well demonstrated in the recent papers proposing the different pan-genome indexes. We implemented multiple reference pan-genome index behaviour using several traditional indexes and BWA read alignment.^4^ With this choice we wanted to show the maximum potential a pan-genome indexing can provide, as BWA is one of the most engineered tools for read alignment.

In a real setting, one should of course use some tailored pan-genome index in order to achieve manageable time and space requirements. We list in Table 3 all existing multiple reference -based pan-genome indexing implementations we are aware of, and their main features. Our main scheme requires a combination of ✓s in columns Model/Set of seqs. and Output alignment/to seq., but such combination is only supported by RLCSA,^8^ whose read alignment support (as discussed in^13^) is not as engineered as that of BWA.^4^ Potentially *Journaled String Tree*^22^ *also has these features, but that approach supports only online* left-to-right search, which may not be scalable for read alignment. As the newer pan-genome indexing proposals^11, 12^ ourperform RLCSA^8^ in practice, and they have a feature to benefit from existing single-reference read aligners such as BWA^4^ in their internal workings, we look forward to their public implementations to be readily usable in combination with our scheme.

**Table 3.**
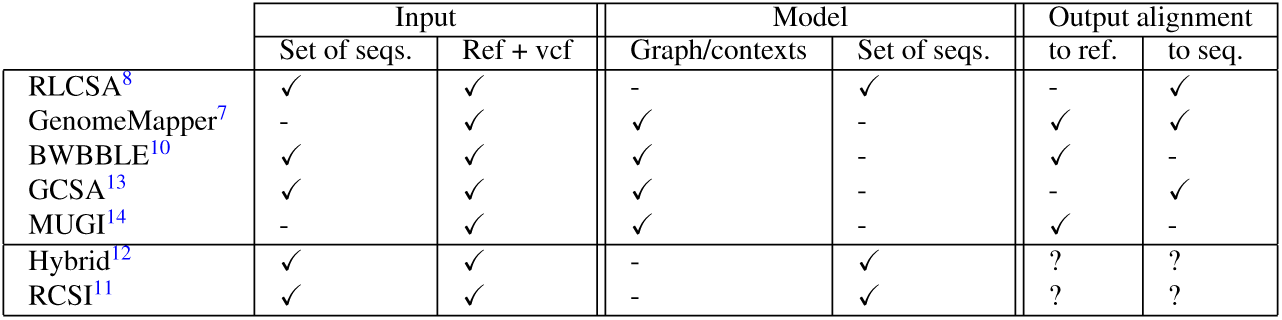
Pan-genome indexing software features. Note that if a software accepts a set of sequences as an input, it is easy to convert a reference plus a set of variants into a set of sequences. Moreover, some tools include a script to do so. However, the reverse transformation is not straightforward. The two last tools described do not have the code publicly available, therefore we are not sure about the output format. The other properties were inferred from the papers.^11, 12^

We note that although we do not currently provide a scalable instance of our main scheme, we did alter a true graph-based pan-genome indexing approach^13^ to provide the same functionality. This provides one scalable and readily usable instance of our approach, and hopefully our experimental results encourage the developers of other pan-genome indexes to look for an output compatible with our approach as well.

Finally, in addition to variation calling, our methods could be extended to other applications such as to support haplotype analysis as in;^15^ see Methods section on the inference of two *ad hoc* references to model a diploid genome with unphased pair of haplotypes.

## Methods

Our overall scheme is illustrated in Fig. 1 on page 2, right. In the following we provide a detailed description of the components of our workflow. We assume the reader is familiar with the acronyms such as BAM and VCF referring to the common file formats used in communicating the read alignments and variation calling results, respectively. We use these to make concrete links to existing workflows in the field. Also, to simplify the exposition, we assume a reference genome is represented as one linear sequence; in practice one can consider a suitably concatenated set of chromosomes.

Our scheme is designed to be modular, and to be used as a preprocessor to any variation calling workflow. The main component is the preprocessor; it takes the raw reads of the donor as an input, and with the use of the pan-genome reference, generates the ad hoc reference.

Then, we resort to the traditional variant calling workflow, VC workflow in our scheme.

Finally, we need to normalize our variants: The VC workflow will call the variants from the ad hoc reference. The normalizer component uses metadata generated from the preprocessor to project the variants to the standard reference.

### Pan-genome preprocessor

The main role of the pan-genome preprocessor is to extract an *ad hoc* reference sequence from the pan-genome using the reads from the donor as an input. An schematic view of the preprocessor is shown in Fig. 2

**Figure 2.**
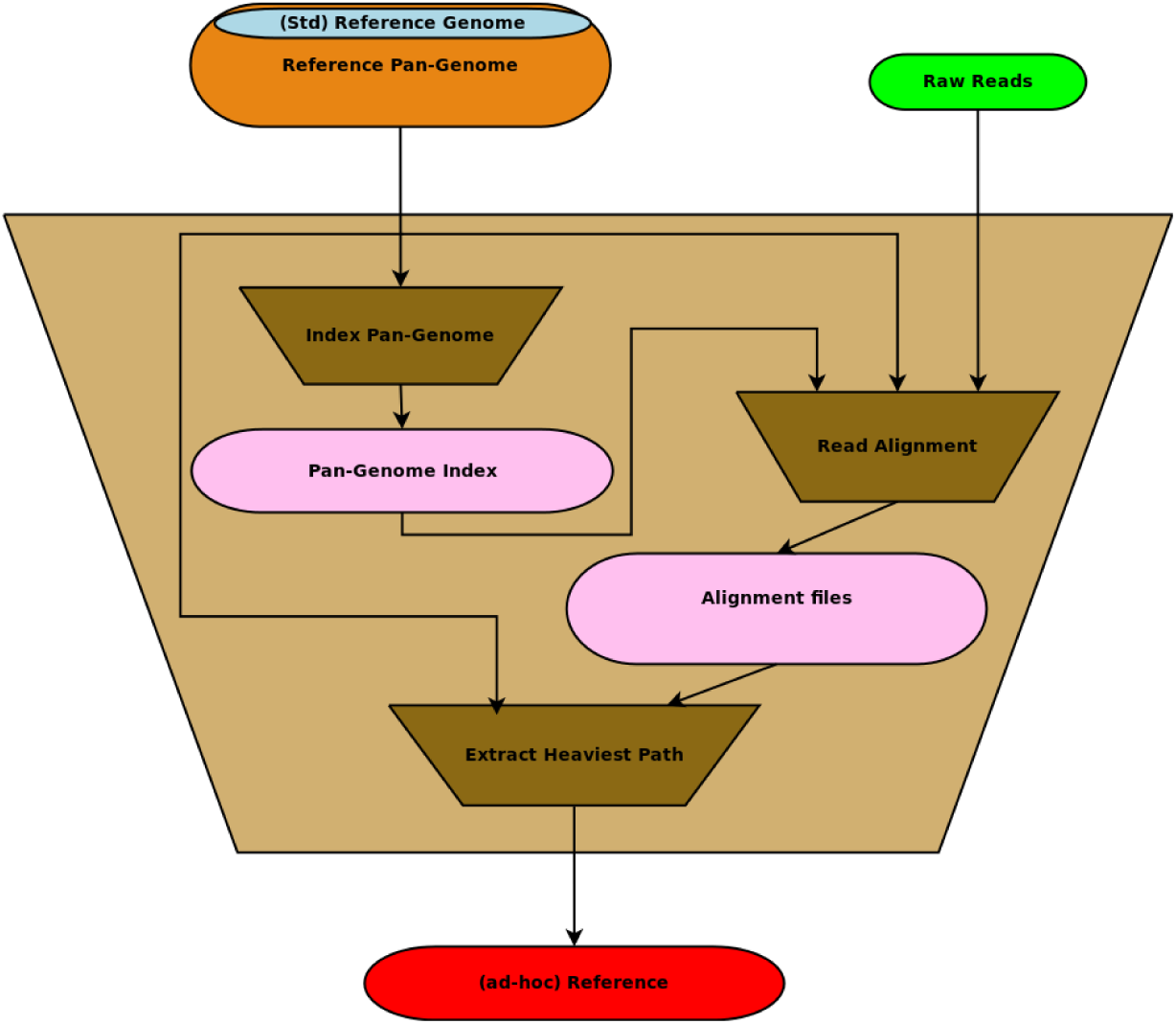
The algorithmic core of the preprocessor is the pan-genome index which allows to map the reads to the pan-genome efficiently. We need to obtain detailed information about the mapped reads (such as BAM files for each sequence in the pan-genome) to enable the extraction of the heaviest path. The heaviest path in the pan-genome is then used as an *ad hoc* reference.

### Pan-genome indexing and read alignment

Following the literature review provided in the Introduction section, the existing pan-genome indexing approaches for read alignment could be classified as follows. There are approaches that consider the input as a set of sequences, those that build a graph or an automata that models the population, and those that consider the specific case of a reference sequence plus a set of variations. However, the boundaries between these categories is not strict, as a set of sequences could be interpreted as a multiple sequence alignment, which in turn can can be turned into a graph.

Our scheme can work with any kind of pan-genome index, provided the following functionality: The output of a read alignment must provide precise information of the alignment: in which of the input sequences the read is mapped, and to which positions. If the index uses an abstract representation of the pan-genome (such as a graph), we need it to be able to provide this information. That is, the alignment needs to be mapped back to the input that was used to build the map. For our scheme we assume that the pan-genome index generates one BAM file for each sequence that conforms the pan-genome. (In the end of this section we leverage these assumptions, and detail how our scheme should be altered when given alignments to a graph-based index.)

Unfortunately, there is currently no off-the-shelf implementation of a pan-genome index that would offer directly our input. To solve this issue (in order to demonstrate our overall scheme) we implemented the following strategy: We consider our input as a multiple sequence alignment, and we store a bitvector that signals all the positions where there is a gap. We index each underlying sequence individually using BWA,^4^ and using the bitvector we are able to efficiently map the occurrences from the underlying sequences to the multiple sequence alignment. This approach does not offer a scalable pan-genome indexing solution, but it provides as good *accuracy* as one can expect a true pan-genome indexing solution to provide. In Discussion section we review the state-of-the-art of current pan-genome indexes and their compatibility with our scheme.

### Heaviest path extraction

After aligning all the reads to the multiple sequence alignment, we extract a recombined (virtual) genome favoring the positions where most reads were aligned.

To do so we propose a generic approach to extract such a heaviest path on a multiple sequence alignment. We define a *score matrix* that has the same dimensions as the multiple sequence alignment representation of the pan-genome. All the values of the score matrix are initially set to 0.

We use the pan-genome index to find the places where each read aligns to the pan-genome. When a read of length *m* is aligned with sequence *i* at position *j*, we increment the scores in *S*[*i*][*j*], *S*[*i*][*j* + 1] *…S*[*i*][*j* + *m–*1]. When all the reads have been processed we have recorded in *S* that the areas with highest scores are those where more reads were aligned. Figures 3 and 4 show a schematic example. (For simplicity of exposition, our description and examples here cover only the case of alignment without indels, but the approach is easy to extend for indels by incrementing scores over the whole aligned region.)

**Figure 3.**
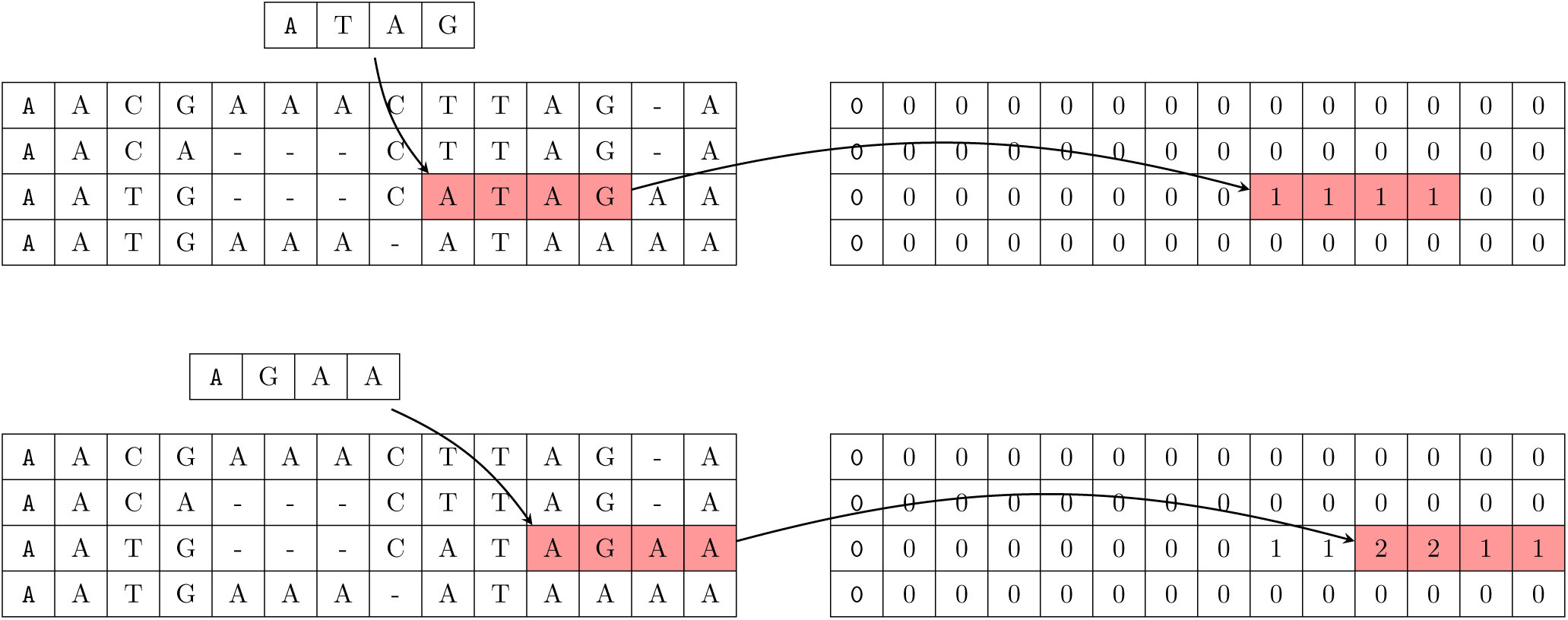
On the left, an example alignment of the reference sequences. On the right the score matrix with the initial values in 0. In the example the first read is “ATAG”, which is aligned within the second sequence at the third position. Then, the corresponding values in the score matrix are incremented by one

**Figure 4.**
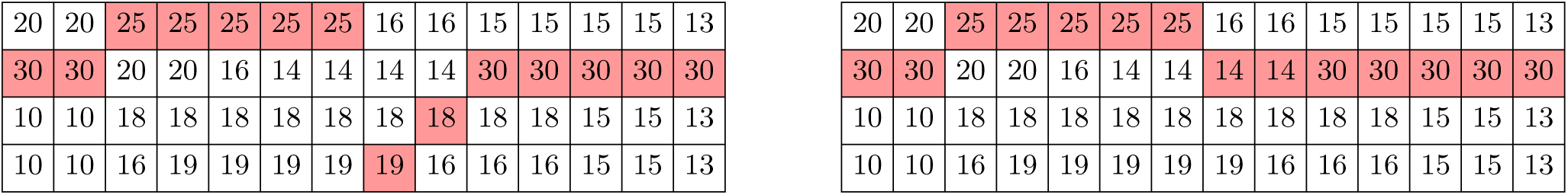
Heaviest path in the score matrix. On the left, without constrains. On the right, the heaviest path is constrained to change two times between sequences. The *ad hoc* references extracted from the pan-genome reference are “AACGAAAATAGA” and “AACGAAACTTAGA” respectively.

To obtain the predicted sequence we consider the matrix as a graph: each cell (*i, j*) represents a node, and for each node (*i, j*) there are *N* outgoing edges to nodes (*i* + 1, *k*), *k* ∈{1, *…, N*}. We add an extra node *A* with *N* outgoing edges to the nodes (1, *k*), and another node *B* with *N* ingoing edges from nodes (*L, k*). To obtain the predicted sequence we use the heaviest weighted path from *A* to *B*. This path is obtained simply by picking the heaviest node at each column of the score matrix. The underlying idea of this procedure is to model structural recombinations among the indexed sequences.

The resulting path might contain too many alternations between sequences in order to maximize the weight. This is fine as a solution to an optimization problem, but the recombination process that we try to emulate might not have that many jumps made of very short segments. To include this fact in our model, we implemented a heaviest path algorithm that receives also a parameter which limits the maximum number of changes of sequences. Figure 4 shows the difference among those scenarios.

To solve the constrained heaviest path problem we use the following dynamic programing formulation. We define a table *V* [1 *…L*][1 *…N*][0 *…Z*] initially set to 0. The values *V* [*i*][*j*][*k*] correspond to the weight of the heaviest path up to character *i*, choosing the last character from sequence *j*, that has made exactly *k* changes of sequences so far. The recursion for the general case (*k* > 0, *i* > 1) is as follows: *V*_*i,j,k*_ = *S*_*i, j*_ + *max*{*V*_*i*−1, *j,k*_, *max*_*j*′ ≠ *j*_ *V*_*i*−1, *j*′,*k*−1_}, and the base case for *k* = 0, *i* > 1 is: *V*_*i,j,*0_ = *S*_*i,j*_ + *V*_*i*−1,*j*_, and for *k* = 0, *i* = 1: *V*_1,*k*,0_ = *S*_1, *j*_.

Once the table is fully computed, the weight of the heaviest path with at most *k\** changes is given by *max j*{*V*_*L,j,k\**_}. To reconstruct the path we need to traceback the solution.

### Variant calling

In Fig. 1 (left) we show a simple workflow that performs variant calling. We do not provide any specifics on purpose: variant calling can be in itself a complex workflow, and it might be tailored for specific type of variants (SNVs, Structural Variants), etc. We aim for a modular and flexible workflow, so any workflow can be plugged there. The only difference is that we will feed it the *ad hoc* reference instead of the standard one. In our experiments, we resort to GATK best practices.^17^

### Normalizer

Finally we need to normalize our set of variants. To do so we apply the variants to the *ad hoc* reference, so that we obtain an alignment between the *ad hoc* reference and the predicted sequence. The metadata generated in the preprocessor stage – while extracting the heaviest path – includes an alignment between the standard reference and the *ad hoc* reference. Using those, we can run a linear time algorithm to obtain the alignment between the standard reference and the predicted sequence. From this alignment, we can generate a vcf file that expresses the predicted sequence as a set of variants from the standard reference.

### Modification to graph representation of pan-genome

Our approach as detailed above is tailored to a multiple sequence alignment representation of a pan-genome, but it is relatively easy to modify the approach to work with a graph representation like one in.^13^ There the pan-genome is stored as a vertex-labeled directed acyclic graph (labeled DAG), and reads are aligned to the paths of this labeled DAG. One can hence store for each vertex the number of read alignments spanning it. Then the heaviest path through this labeled DAG is easy to compute by dynamic programming in a topological ordering of the graph: The weight of the heaviest path *h*(*v*) to a vertex *v* is max_*v* ′ *N*−(*v*)_ *h*(*v*′) + *w*(*v*), where *w*(*v*) is the weight of a vertex and *N*^−^(*v*) is the set of vertices connected with a in-coming arc to *v*. connected with a in-Traceback from the sink vertex shows the vertices through which a heaviest path goes, and one can output the corresponding vertex labels as the *ad hoc* reference.

The difference to the multiple alignment heaviest path is that the number of recombinations cannot be limited when using the graph representation.

Another part that is different is the normalizer module to map the variants predicted from the *ad hoc* reference to the standard reference. For this, the original proposal in^13^ already records the path spelling the standard reference, so while extracting the heaviest path one can detect the intersection to the standard reference path and store the corresponding projection as an alignment.

### Inference of pair of haplotypes

For variation calling, the inference of a single representative haplotype sequence is a convenience. However, to get a better prediction of the underlying diploid genome one can modify the heaviest path algorithms to produce two predictions. One way to do this is to remove the coverages along the path of the first *ad hoc* reference and run the heaviest path algorithm again to produce a second *ad hoc* reference. This results into a pair of unphased haplotype sequences. Evaluation methods for such a prediction sheme have been studied in.^23^

## References

1. Burrows, M. & Wheeler, D. A block-sorting lossless data compression algorithm. Tech. Rep. 124, Digital Equipment Corporation (1994).

2. Ferragina, P. & Manzini, G. Opportunistic data structures with applications. In Foundations of Computer Science, 2000. Proceedings. 41st Annual Symposium on, 390–398 (IEEE, 2000).

3. Ferragina, P. & Manzini, G. Indexing compressed text. Journal of the ACM (JACM) 52, 552–581 (2005).

4. Li, H. & Durbin, R. Fast and accurate short read alignment with burrows–wheeler transform. Bioinformatics 25, 1754–1760 (2009).

5. Langmead, B., Trapnell, C., Pop, M., Salzberg, S. L. et al. Ultrafast and memory-efficient alignment of short dna sequences to the human genome. Genome biol 10, R25 (2009).

6. Li, R., Li, Y., Kristiansen, K. & Wang, J. Soap: short oligonucleotide alignment program. Bioinformatics 24, 713–714 (2008).

7. Schneeberger, K. et al. Simultaneous alignment of short reads against multiple genomes. Genome Biology 10, R98 (2009).

8. Mäkinen, V., Navarro, G., Sirén, J. & Välimäki, N. Storage and retrieval of highly repetitive sequence collections. Journal of Computational Biology 17, 281–308 (2010).

9. Deorowicz, S., Danek, A. & Grabowski, S. Genome compression: a novel approach for large collections. Bioinformatics 29, 2572–2578 (2013).

10. Huang, L., Popic, V. & Batzoglou, S. Short read alignment with populations of genomes. Bioinformatics 29, 361–370 (2013).

11. Wandelt, S., Starlinger, J., Bux, M. & Leser, U. Rcsi: Scalable similarity search in thousand (s) of genomes. Proceedings of the VLDB Endowment 6, 1534–1545 (2013).

12. Ferrada, H., Gagie, T., Hirvola, T. & Puglisi, S. J. Hybrid indexes for repetitive datasets. Philosophical Transactions of the Royal Society A 372(2014).

13. Sirén, J., Välimäki, N. & Mäkinen, V. Indexing graphs for path queries with applications in genome research. IEEE/ACM Transactions on Computational Biology and Bioinformatics 11, 375–388 (2014).

14. Danek, A., Deorowicz, S. & Grabowski, S. Indexing large genome collections on a pc. PLoS ONE 9, e109384 (2014).

15. Dilthey, A., Cox, C., Iqbal, Z., Nelson, M. R. & McVean, G. Improved genome inference in the mhc using a population reference graph. Nature Genetics 47, 682–688 (2015).

16. Wandelt, S. & Leser, U. Mrcsi: Compressing and searching string collections with multiple references. PVLDB 8, 461–472 (2015).

17. Auwera, G. A. et al. From fastq data to high-confidence variant calls: the genome analysis toolkit best practices Current Protocols in Bioinformatics 11–10 (2013).

18. Li, H. A statistical framework for snp calling, mutation discovery, association mapping and population genetical parameter estimation from sequencing data. Bioinformatics 27, 2987–2993 (2011).

19. Wittler, R., Marschall, T., Schönhuth, A., & Mäkinen, V. Repeat- and error-aware comparison of deletions. Bioinformatics (2015). In press, doi:10.1093/bioinformatics/btv304.

20. Horton, R. et al. Variation analysis and gene annotation of eight MHC haplotypes: The MHC haplotype project. Immunogenetics 60(2007).

21. Khurana, E. et al. Integrative annotation of variants from 1092 humans: Application to cancer genomics. Science 342 (2013).

22. Rahn, R., Weese, D. & Reinert, K. Journaled string tree – a scalable data structure for analyzing thousands of similar genomes on your laptop. Bioinformatics 30, 3499–3505 (2014).

23. Mäkinen, V. & Valenzuela, D. Recombination-aware alignment of diploid individuals. BMC Genomics 15, S15 (2014).

